# Brain Age in Conduct Disorder: A Mega-Analysis of the ENIGMA Antisocial Behavior Working Group

**DOI:** 10.64898/2026.01.20.700567

**Authors:** Jules R. Dugré, Yidian Gao, Marlene Staginnus, Abigail A. Marsh, Alexandra Kypta-Vivanco, Ana I. Cubillo, Andrea Dietrich, Anka Bernhard, Anne Martinelli, Anouk H. Dykstra, Areti Smaragdi, Arjun Sethi, Arun L.W. Bokde, Barbara Franke, Celso Arango, Charlotte P.S. Boateng, Christina Stadler, Christine M. Freitag, Christopher D. Townsend, Christopher S. Monk, Cindy C Hagan, Colter Mitchell, Daifeng Dong, Dana E. Díaz, Daniel Brandeis, Declan Murphy, Denis G. Sukhodolsky, Edmund J.S. Sonuga-Barke, Elise M. Cardinale, Essi Viding, Gregor Kohls, Gunter Schumann, Harriet Cornwell, Harriet Phillips, Heidi B. Westerman, Hui Chen, Ilyas Sagar-Ouriaghli, Jack C. Rogers, Jan K. Buitelaar, Jean-Luc Martinot, Jeffrey C. Glennon, Jiansong Zhou, Jibiao Zhang, Jilly Naaijen, Josefina Castro-Fornieles, Kalina J. Michalska, Karen Gonzalez-Madruga, Karim Ibrahim, Kate Sully, Kathryn Berluti, Kerstin Konrad, Kim Lamers, Leah E. Mycue, Luca Passamonti, Luke W. Hyde, Maaike Oosterling, Maria Jose Penzol Alonso, Michael C Stevens, Michael Craig, Mireia Rosa-Justicia, Montana L. Ploe, Nathalie Holz, Nic J.A. van der Wee, Nicola Toschi, Nimrah Jabeen, Nora M. Raschle, Olivier F. Colins, Paramala Santosh, Pascal-M Aggensteiner, Pieter J Hoekstra, Qingsen Ming, Qiong Wu, Rebecca L. Jackson, Ren Ma, Robert J.R. Blair, Robert Vermeiren, Ruth Pauli, Ruth Roberts, S. Alexandra Burt, Sahil Bajaj, Sally C. Chester, Silvia Minosse, Sylvane Desrivieres, Tobias Banaschewski, Ulrike M.E. Schulze, Xianliang Chen, Xiaoping Wang, Xiaoqiang Sun, Yali Jiang, Neda Jahanshad, Sophia I. Thomopoulos, Christopher R.K. Ching, Melody J.Y. Kang, Paul M. Thompson, Daniel S. Pine, Arielle Baskin-Sommers, Charlotte A.M. Cecil, Moji Aghajani, Graeme Fairchild, ENIGMA Antisocial Behavior Consortium, Stephane A. De Brito

## Abstract

Conduct disorder (CD) is the leading global cause of mental health burden in children and adolescents and has recently been hypothesized to be a neurodevelopmental disorder. Although prior research has identified neuroanatomical differences associated with CD, it remains unclear whether these differences reflect atypical brain development. Here, we investigated the difference between an individual’s brain age and chronological age as a proxy for variations in brain maturation. Using a pretrained model, we estimated brain age from structural neuroimaging data obtained from 1,119 youth with CD and 1,183 typically developing controls across 14 international cohorts participating in the ENIGMA-Antisocial Behavior Working Group. Youth with CD exhibited a statistically robust but small acceleration in brain age compared to typically developing youth (around 0.50 years), which was restricted to the adolescence-onset subtype of the disorder. Our large-scale, coordinated analysis provides the first evidence of accelerated neurodevelopment as a potential mechanism underlying CD.

## Introduction

Conduct Disorder (CD) is characterized by a persistent pattern of behavior that violates societal norms and the rights and welfare of others^1^. It is typically diagnosed in youth^1^, defined here as a developmental stage extending through the mid-20s^2^. Youth with CD represent a highly heterogeneous group, with substantial inter-individual variability in clinical presentation largely influenced by factors such as age of onset (i.e., childhood-onset [<10 years] *vs*. adolescent-onset [≥10 years]) and the presence of limited prosocial emotions (e.g., lack of remorse or guilt, callousness or lack of empathy)^1^, often referred to as callous-unemotional traits ^3^. Across the globe, CD associated with the highest burden of any mental disorder in children and adolescent aged 0–14 years^4^, and imposes significant economic and societal costs^5,6^. Moreover, CD has been prospectively linked to many other mental illnesses, as well as poorer psychosocial adjustment and health outcomes in adulthood^7^. Given its widespread impact, research investigating putative developmental processes should be a priority.

Growing evidence suggests that CD may be a neurodevelopmental disorder^7–9^, especially in youth whose symptoms emerge during childhood^10^. For instance, it shares genetic influences^11^ and neurobiological features^12,13^ with other neurodevelopmental disorders, particularly attention-deficit/hyperactivity disorder. Furthermore, prior meta-analyses have shown that youth with CD exhibit grey matter volume reductions in brain regions that undergo substantial development during childhood and adolescence (e.g., amygdala, anterior insula/ventrolateral prefrontal cortices)^14,15^. Of note, the largest neuroimaging study to date on CD, a collaborative effort of the ENIGMA-Antisocial Behavior Working Group (ENIGMA-ASB), demonstrated that youth with CD exhibit robust and widespread reductions in cortical surface area and subcortical volumes (i.e., thalamus, amygdala, hippocampus, nucleus accumbens), but few differences in cortical thickness^16^. Despite considerable progress in characterizing structural brain alterations in youth with CD, the neurodevelopmental processes driving these differences remain poorly understood.

In the past decade, researchers have developed a new multivariate method to quantify whether an individual’s structural brain metrics reflect accelerated or decelerated brain maturation^17^. This is typically achieved by predicting chronological age from neuroimaging brain metrics (e.g., surface area, cortical thickness, subcortical volume). The resulting difference between predicted and actual age -referred to as the brain predicted-age difference (brain-PAD) - is now widely used to capture deviations from normative brain maturation^17^. A brain-PAD significantly above zero would thus reflect accelerated brain age (brain age > chronological age), while a brain-PAD significantly below zero would indicate decelerated brain age (brain age < chronological age). This approach has opened promising avenues for understanding psychiatric disorders.

For instance, accelerated brain aging has been commonly observed in adults with psychiatric^18–24^ and neurological disorders^25^ compared to controls. In children, a preliminary study showed decelerated brain maturation in youth with attention-deficit/hyperactivity disorder^26^, although a recent mega-analysis of multiple cohorts revealed no signs of such negative brain age gap in children with neurodevelopmental disorders (autism spectrum disorder and attention-deficit hyperactivity disorder)^27^. Nevertheless, young adults with a history of conduct problems^28^ have shown an average brain-PAD below zero, indicative of a potentially decelerated brain maturation in youth with externalizing problems^29^. Another study found no significant group differences in brain-PAD between adult offenders with a history of violence and healthy controls, although brain-PAD and psychopathic traits were negatively correlated (*r*=-.31)^30^. While these findings suggest that CD and psychopathic traits may be associated with decelerated brain maturation, they are largely limited to adult males and thus may not be generalizable to youth. Indeed, the use of adult samples relying on retrospective accounts of conduct problems, in the absence of formal CD diagnoses, limits the relevance of prior findings for understanding CD in youth. Furthermore, the extent to which these findings generalize across the sexes remains unknown. As both the clinical features of CD and brain structures undergo marked developmental changes around puberty, examining brain age could provide key insights into its neurodevelopmental mechanisms.

In this preregistered and collaborative effort from the ENIGMA-ASB Working Group (https://doi.org/10.17605/OSF.IO/U4D37), we tested whether youth with CD show differences in brain-PAD compared to typically developing youth. First, based on the available evidence on the association between externalizing problems and brain aging^26,28–30^, as well as recent longitudinal studies of brain development^31–33^, we hypothesized that youth with CD would display decelerated brain age relative to their chronological age (brain-PAD significantly below zero). Second, we hypothesized that brain-PAD would negatively correlate with dimensionally assessed conduct problems in youth with CD. Third, we expected that decelerated brain age would be more pronounced in the childhood-onset CD subtype than in typically developing youth and adolescent-onset CD, consistent with neurodevelopmental impairments specific to the childhood-onset subgroup^10^. Moreover, because CD subgroups with high callous-unemotional traits often exhibit earlier onset and a more severe clinical presentation than those with low traits^7^, we hypothesized that the high-trait subgroup would show the greatest deceleration in brain age, relative to the low-trait subgroup and typically developing youth. We also examined whether sex moderated the association between CD and brain age. Although previous work has shown that the association between brain age and history of CP was limited to males^28^, we did not formulate specific hypotheses on this due to limited evidence in CD. Finally, we expected that decelerated brain age would generalize to non-overlapping participants with elevated conduct problems but without a formal diagnosis of CD derived from 10 ENIGMA-ASB cohorts (926 subthreshold CD youth *vs.* 922 controls).

## Results

### Sample Characteristics

In the current study, we included 1,119 youth with CD (328 females, 791 males; mean age of 13.7, SD = 3.04) and 1183 typically developing youth (436 females, 747 males; mean age of 13.3, SD=3.03) from the 14 participating cohorts of the ENIGMA-ASB Working Group. See Table 1 for demographic and clinical characteristics of the sample.

**Table 1.** Demographic and clinical characteristics of the included cohorts in the main analysis on youth with Conduct Disorder

| Cohorts | N | Age range, yrs | Typically developing youth |  |  |  | Youth with Conduct Disorder |  |  |  | Conduct disorder subgroups |  |  |  |
| --- | --- | --- | --- | --- | --- | --- | --- | --- | --- | --- | --- | --- | --- | --- |
|  |  |  | n | F:M | Age, yrs | IQ | n | F:M | Age, yrs | IQ | CO-CD | AO-CD | CU- | CU+ |
| ABCD (3.0, baseline)*† | 572 | 9–10 | 286 | 81:205 | 9.5 (0.50) | 94.9 (15.3) | 286 | 85:201 | 9.5 (0.50) | 94.6 (16.4) | 256 | 30 | - | - |
| BESD | 86 | 14–19 | 36 | 0:36 | 16.7 (1.32) | 97.3 (9.41) | 50 | 0:50 | 16.4 (1.34) | 95.9 (6.40) | - | - | 38 | 12 |
| Boys Town | 349 | 10–19 | 169 | 62:107 | 13.7 (2.44) | 108.8 (12.56) | 180 | 62:118 | 15.3 (1.71) | 99.3 (11.62) | - | - | 64 | 112 |
| Cambridge Female | 46 | 14–19 | 23 | 23:0 | 17.0 (0.88) | 105.7 (9.34) | 23 | 23:0 | 16.7 (1.66) | 99.5 (8.11) | 5 | 17 | 11 | 8 |
| Cambridge Male | 88 | 16–21 | 25 | 0:25 | 18.0 (1.08) | 101.6 (9.10) | 63 | 0:63 | 17.2 (1.11) | 98.8 (8.57) | 26 | 37 | 24 | 26 |
| CD-Zhou | 27 | 16–18 | 13 | 0:13 | 16.9 (0.28) | - | 14 | 0:14 | 17.0 (0.56) | - | - | - | - | - |
| CDKid | 31 | 12–19 | 13 | 0:13 | 15.7 (2.02) | 109.8 (11.06) | 18 | 0:18 | 15.4 (1.54) | 98.7 (7.14) | 12 | 6 | - | - |
| CSU-Yao | 145 | 12–17 | 71 | 0:71 | 15.5 (0.56) | 108.9 (9.03) | 74 | 0:74 | 14.5 (1.14) | 100.8 (12.96) | 5 | 64 | - | - |
| FemNAT-CD* | 612 | 9–18 | 365 | 217:148 | 14.3 (2.53) | 103.6 (11.56) | 247 | 118:129 | 14.6 (2.16) | 95.1 (12.48) | 124 | 104 | 81 | 153 |
| IMAGEN (baseline)*† | 126 | 13–15 | 63 | 24:39 | 14.0 (0.46) | - | 63 | 24:39 | 14.0 (0.46) | - | - | - | - | - |
| K23 | 34 | 12–18 | 24 | 14:10 | 14.8 (1.82) | - | 10 | 4:6 | 15.9 (0.99) | - | - | - | 5 | 1 |
| MATRICS/Aggressotype*† | 93 | 7–18 | 49 | 7:42 | 12.8 (2.63) | 100.6 (10.91) | 44 | 6:38 | 13.1 (2.82) | 98.9 (10.93) | - | - | 20 | 16 |
| Southampton Family Study | 71 | 13–18 | 35 | 6:29 | 16.0 (1.36) | 104.2 (10.03) | 36 | 4:32 | 16.1 (1.41) | 93.6 (11.42) | 20 | 16 | 18 | 18 |
| Yale† | 22 | 9–16 | 11 | 2:9 | 11.7 (1.62) | 103.3 (14.40) | 11 | 2:9 | 12.2 (2.27) | 102.1 (12.98) | - | - | 2 | 9 |
| Total (14 samples) | 2302 | 7–21 | 1183 | 436:747 | 13.3 (3.03) | 102.2 (13.47) | 1119 | 328:791 | 13.7 (3.04) | 96.7 (13.13) | 448 | 274 | 263 | 355 |

*Note.* The reported values reflect n or mean (SD), unless otherwise indicated. N refers to the total number of participants from a specific cohort that were included in the current study. Information on sex and age was available for all participants, whereas IQ, age-of-onset status, and CU traits data were not always available. F=Females. M=Males. IQ=intelligence quotient. CO-CD = Childhood-Onset Conduct Disorder; AO-CD = Adolescent-Onset Conduct Disorder; CU=callous-unemotional (- = Low; + = High).
\* Multi-site or multi-scanner sample.
† Control group matched on age and sex (and IQ, if available) using propensity score matching;

### Brain Age in Conduct Disorder

The BrainAGE Global model showed adequate generalization in our sample, as demonstrated by a strong correlation (*r* = 0.81) between predicted age from neuroanatomical features and chronological age, and a relatively low mean absolute error (2.71 years). The model demonstrated similar performance for CD and typically developing youth, as well as for males and females (see Supplementary Fig. 1). Model performance was slightly lower for adolescent-onset CD subtype (r =.61, MAE = 3.60), compared to childhood-onset CD subtype (r = .85, MAE = 2.25) and typically developing (r = .79, MAE =2.56). Similarly, CD subtype with high callous-unemotional traits showed slightly lower performance metrics (r = .54, MAE = 3.39) compared to those with low callous-unemotional traits (r = .63, MAE = 3.19) and typically developing groups (r = .69, MAE = 2.88).

Contrary to our predictions, youth with CD showed a significantly greater positive brain-PAD compared to typically developing youth (*p* = 0.0002, Cohen’s *d* = 0.14, 95% CI: 0.05–0.22) after adjusting for sex, chronological age, and chronological age^2^, with cohorts included as a random intercept to account for between-site variability. (Table 2, Fig. 1A). The mean brain-PAD was significantly above zero (+0.45 years), indicating greater brain age compared to chronological age. A jackknife resampling procedure, sequentially excluding each cohort, confirmed the stability of the observed effect across cohorts (Mean estimate = 0.456, SE = 0.14, 95% CI: 0.18 - 0.73). All iterations were statistically significant (*p*-values ranging from 0.00003 to 0.0018), suggesting that the observed accelerated brain age was not driven by any specific cohort (see Supplementary Fig 2). The group difference in brain age was statistically robust after accounting for group differences in intelligence quotient (1016 CD *vs*. 1078 typically developing participants, *p* = 0.00048, *d* = 0.14), attention-deficit/hyperactivity disorder comorbidity (1102 CD *vs*. 1181 typically developing*, p* = 0.0086, *d* = 0.09), and medication status (1002 CD *vs*. 1051 typically developing*, p* = 0.0014, *d* = 0.13). No interactions with chronological age were found - there was no two-way (diagnosis-by-chronological age, *p* = 0.992) or three-way (diagnosis-by- chronological age-by-sex, *p* = 0.745) interaction with age. There was also no diagnosis-by-sex interaction (*p* = 0.868). Dimensionally assessed conduct problems were not associated with brain-PAD within the CD group (n = 617, *p* = 0.348, *r* = 0.03).

**Fig. 1.**
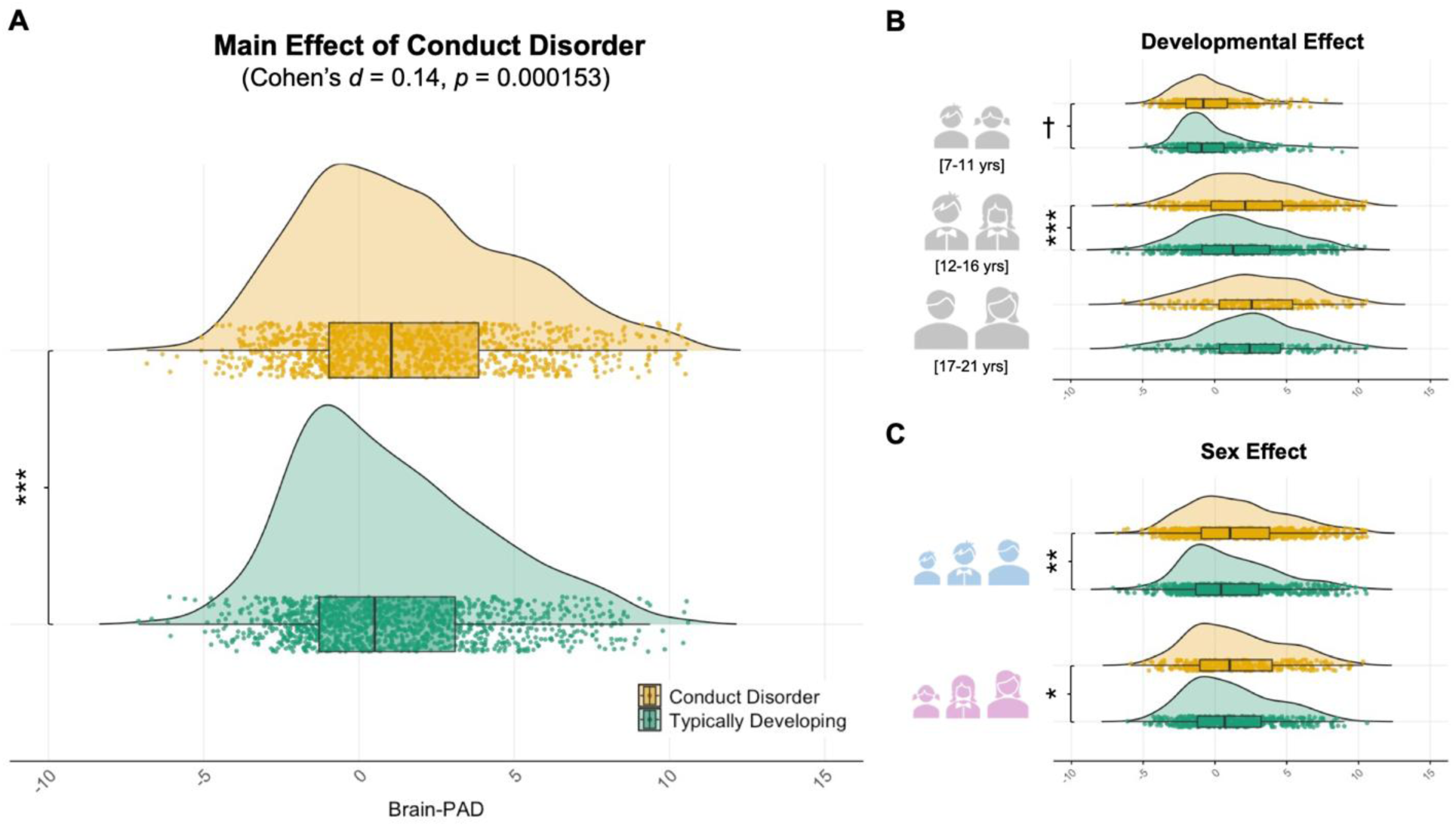
Case-Control comparison of Brain-PAD between youth with CD and Typically Developing youth. **A. A** linear mixed effect model showed that youth with CD exhibited an accelerated brain age compared with typically developing youth (*b* = 0.45, *p* = 0.000153), representing a Cohen’s d effect size of 0.14 after adjusting for chronological age, chronological age^2^, sex, and cohort. **B.** Sensitivity analyses examined whether this effect was consistent across developmental periods approximating pubertal stages, which are childhood (7-11 years old, n = 742), adolescence (12-16 years old, n = 1129), and late adolescence/young adulthood (17-21, n = 431). Results showed strongest accelerated brain age in early to mid-adolescence (*d* = 0.17, *p* = 0.001), and weakest in young adulthood (*d* = 0.04, *p* = 0.635), with intermediate effect observed in childhood (*d* = 0.12, *p* = 0.065). **C.** Sensitivity analyses were also performed to assess this effect was consistent across sexes, and revealed that the accelerated brain age was observed across males (*n* = 1,538, *d* = 0.13, *p* = 0.0019), and females (*n* = 764, *d* = 0.14, *p* = 0.04). (^†^ *p* = 0.065; * *p* < 0.05; ** *p* < 0.01; *** *p* < 0.001).

**Table 2.** Chronological Age, Brain Age, and Brain Predicted Age Differences per cohorts.

| Cohorts | Typically Developing Youth |  |  |  | Youth with Conduct Disorder |  |  |  |
| --- | --- | --- | --- | --- | --- | --- | --- | --- |
|  | n | Chronological age | Brain Age | Brain-PAD | n | Chronological age | Brain Age | Brain-PAD |
| ABCD (3.0, baseline)*† | 286 | 9.5 (0.50) | 8.6 (1.83) | -0.95 (1.78) | 286 | 9.5 (0.50) | 8.79 (2.16) | -0.66 (2.09) |
| BESD | 36 | 16.7 (1.32) | 17.8 (3.13) | 1.12 (2.53) | 50 | 16.4 (1.34) | 19.2 (3.58) | 2.75 (3.27) |
| Boys Town | 169 | 13.7 (2.44) | 15.2 (3.98) | 1.45 (2.94) | 180 | 15.3 (1.71) | 17.7 (3.95) | 2.43 (3.42) |
| Cambridge Female | 23 | 17.0 (0.88) | 19.0 (2.80) | 1.94 (2.85) | 23 | 16.7 (1.66) | 18.5 (2.88) | 1.72 (3.00) |
| Cambridge Male | 25 | 18.0 (1.08) | 19.3 (3.62) | 1.28 (3.39) | 63 | 17.2 (1.11) | 18.7 (3.59) | 1.45 (3.28) |
| CD-Zhou | 13 | 16.9 (0.28) | 22.6 (2.50) | 5.69 (2.55) | 14 | 17.0 (0.56) | 23.6 (2.50) | 6.61 (2.19) |
| CDKid | 13 | 15.7 (2.02) | 19.3 (3.87) | 3.64 (2.78) | 18 | 15.4 (1.54) | 18.1 (3.24) | 2.70 (3.33) |
| CSU-Yao | 71 | 15.5 (0.56) | 18.9 (2.15) | 3.42 (2.22) | 74 | 14.5 (1.14) | 18.6 (2.81) | 4.09 (2.88) |
| FemNAT-CD* | 365 | 14.3 (2.53) | 15.9 (4.36) | 1.55 (3.25) | 247 | 14.6 (2.16) | 16.4 (4.03) | 1.73 (3.34) |
| IMAGEN (baseline)*† | 63 | 14.0 (0.46) | 13.9 (1.33) | -0.07 (1.26) | 63 | 14.0 (0.46) | 14.1 (1.54) | 0.10 (1.44) |
| K23 | 24 | 14.8 (1.82) | 14.9 (5.68) | 0.10 (4.45) | 10 | 15.9 (0.99) | 17.8 (3.87) | 1.90 (3.43) |
| MATRICES/Aggressotype*<br>† | 49 | 12.8 (2.63) | 13.5 (4.60) | 0.77 (3.10) | 44 | 13.1 (2.82) | 15.4 (5.13) | 2.37 (3.69) |
| Southampton Family Study | 35 | 16.0 (1.36) | 21.3 (3.77) | 5.30 (3.44) | 36 | 16.1 (1.41) | 21.7 (3.50) | 5.59 (2.71) |
| Yale† | 11 | 11.7 (1.62) | 12.9 (4.24) | 1.14 (2.97) | 11 | 12.2 (2.27) | 13.0 (4.70) | 0.81 (2.80) |
| Total (14 samples) | 1183 | 13.3 (3.03) | 14.4 (5.12) | 1.06 (3.14) | 1119 | 13.7 (3.04) | 15.2 (5.30) | 1.54 (3.37) |

*Note.* Brain Age and Brain-PAD were computed using BrainAGE Global model of CentileBrain. Brain Age refers to Neuroimaging-Predicted Age. Brain-PAD = Brain-Predicted Age Difference ( $\Delta$ brain age - chronological age)

Subsequent analyses explored whether the greater positive brain-PAD in CD compared to typically developing youth was consistent across developmental stages and/or across sexes. As shown in Fig. 1B, the CD group still showed greater positive brain-PAD when restricting the analyses to adolescents aged 12-16 years (568 CD *vs*. 561 typically developing; *p* = 0.001, *d* = 0.17). This effect was, however, not statistically significant among children aged 7 to 11 years (333 CD *vs*. 409 typically developing; *p* = 0.065, *d* = 0.12) or late adolescents/young adults aged 17 to 21 years old (218 CD *vs*. 213 typically developing; *p* = 0.635, *d* = 0.04), though the effects in these developmental stages were in the same direction and in close to significance levels. Moreover, accelerated brain age was also found when restricting the analyses to males (*n* = 1,538, *p* = 0.0019, *d* = 0.13), or females (*n* = 764, *p* = 0.04, *d* = 0.14), with very similar effect sizes in each sex (Fig. 1C).

### Impact of Callous-Unemotional Traits and Age-of-Onset of CD

To better understand accelerated brain age, considering the substantial heterogeneity among youth with CD, we conducted sub-analyses to test the generalizability of this effect across subgroups. Table 1 outlines which cohorts contributed to these analyses, with further details provided in Supplementary Methods.

First, we examined the differences in brain-PAD across subgroups of CD by subdividing them based on whether they had elevated callous-unemotional traits, based on the normative cut-off for the ICU. Data on CU traits was available for 618 youth with CD (9 out of 14 cohorts). We observed greater brain-PAD in those with high levels of callous-unemotional traits compared to the other groups (*p* = 0.008, *d* = 0.15; Fig. 2). Pairwise comparisons revealed that this effect was mainly driven by differences between the high callous-unemotional traits subgroup and typically developing youth (*q* = 0.024, *d* = .16), but not those with low levels of callous-unemotional traits (*q* = 0.474, *d* = 0.05). Those with low levels of callous-unemotional traits did not significantly differ from typically developing youth (*q* = 0.107, *d* = 0.12), although the effect was in the direction of accelerated brain age. Using a dimensional approach, there was no association between callous-unemotional traits and brain-PAD, either focusing on youth with CD (*n* = 618, *r* = 0.03, *p* = 0.416) or across the full sample (*n* = 1,342, *r* = 0.04, *p* = 0.088).

**Fig. 2.**
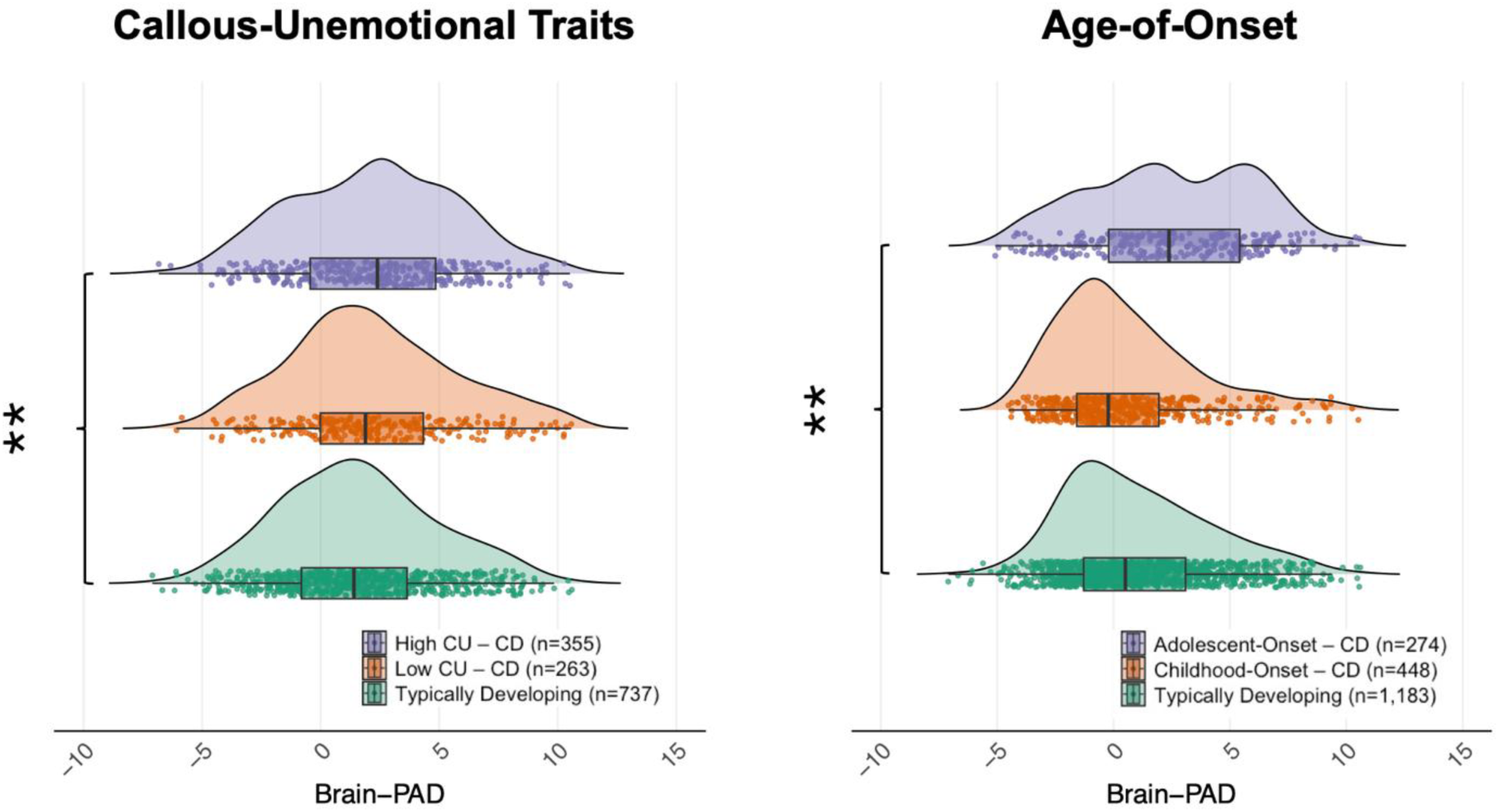
Sub-analyses investigating the impact of Callous-Unemotional (CU) Traits and CD Age-Of-Onset on Brain-Predicted Age Difference (Brain-PAD) differences between the CD and typically developing groups. Brain-PAD refers to the difference between predicted age from neuroanatomical measures and chronological age. Ridge plot on the left panel represents the distributions of Brain-PAD among CD youth with clinical/high (n=355) and low (n=263) levels of CU traits, and typically developing youth (n=737). Ridge plot on the right panel denotes the distributions of Brain-PAD across CD youth with a childhood-onset (n=448) and adolescent-onset (n=274), and the typically developing group (n=1,183). * *p* < 0.05; ** *p* < 0.01; *** *p* < 0.001 without FDR adjustments).

Second, we investigated potential differences in brain-PAD across subgroups of youth with CD based on their age-of-onset (before [n = 448] or after 10 years of age [n = 274], with data available from 7 out of 14 cohorts). We observed a greater brain-PAD in adolescent-onset CD compared to the other groups (*p* = 0.0076, *d* = 0.15; Fig. 2). Pairwise comparisons revealed that this effect was mainly driven by differences between the adolescent-onset CD and typically developing groups (*q* = 0.024, *d* = 0.15), but not between the adolescent-onset and childhood-onset CD subgroups (*q* = 0.139, *d* = 0.11). The childhood-onset CD subgroup did not significantly differ from the typically developing group in brain-PAD (*q* = 0.529, *d* = 0.03).

### Brain Age in youth with elevated Conduct Problems

In line with our previous study^16^, we further aimed to examine whether the brain-PAD difference observed in youth with CD would extend to those with elevated conduct problems, which represents a mix of sub-threshold and undiagnosed cases. A total of 926 youth with elevated conduct problems but without a formal diagnosis of CD (405 females, 521 males; mean chronological age of 11.7 years, SD=2.38) and 922 controls (435 females, 487 males; mean chronological age of 11.6 years, SD=2.34) from 10 cohorts of ENIGMA-ASB were pooled. None of the participants included in this analysis overlapped with those from the main analysis. Of note, this sample was younger (M = 11.7) than the sample from the main CD analysis (M = 13.4).

Both youth with elevated conduct problems (-0.50 years, SD = 1.67) and controls (-0.59 years, SD = 1.58) displayed a mean brain-PAD below zero, suggesting decelerated brain age (Supplementary Table 2, Supplementary Fig. 2). However, no significant difference was found between the groups (*b* = 0.076, SE = .073, *p* = 0.299, *d* = .05).

## Discussion

To our knowledge, this study is the first to investigate brain age in youth with CD in comparison with TD youth. Building on prior work conducted on relatively small samples, we leveraged data from 14 international cohorts, comprising an unprecedentedly large sample of 1,119 youth with CD and 1,183 typically developing youth. We first hypothesized that youth with CD would exhibit decelerated brain age compared to their typically developing counterparts. Contrary to our expectations, analyses revealed that youth with CD showed accelerated brain age relative to typically developing youth, which was not driven by any specific cohort. Furthermore, this effect was similarly observed across sexes but appeared limited early- to mid-adolescence (ages 12-16). Again, in contrast to our hypothesis, both CD subtype with elevated callous-unemotional traits and adolescent-onset CD subtype showed accelerated brain age compared to typically developing youth, although they did not significantly differ from CD subtype with low callous-unemotional traits and childhood-onset CD subtype, respectively. Neither CD subtype with low callous-unemotional traits nor childhood-onset CD subtype significantly differed from typically developing youth. Finally, we found that this accelerated brain age did not generalize to an independent sample of youth with subthreshold CD symptoms, contrary to our expectations. Taken together, our study provides evidence that accelerated brain age may be a potential neurodevelopmental mechanism underpinning CD.

### CD Diagnosis relates to Brain Age

We show for the first time that youth with CD are characterized by accelerated brain age (+0.45 years), with an effect size approximately twice as large as those reported in neurodevelopmental disorders which were unrelated to brain age gap^27^. This accelerated brain age co-occurs with widespread reductions in cortical surface area and subcortical volume previously reported in the ENIGMA-ASB youth sample^16^, which provides further support for a neurodevelopmental conceptualization of CD^8,9^. In contrast to our findings, two previous studies reported a negative association between brain age and externalizing symptoms. This apparent discrepancy is likely attributable to differences in sample size, developmental timing (youth vs. adulthood), and study settings (clinical vs. community). Although delayed brain maturation has been proposed as a characteristic of youth with externalizing problems in developmental models^29^, some studies have failed to support this hypothesis in youth with CD^34^, while others have found that the severity of conduct problems was instead associated with accelerated cortical thinning, particularly during late adolescence^35^.

Accelerated brain age in youth has been associated with many environmental factors, such as adverse childhood experiences and neighborhood disadvantage^36,37^. As many youth with CD experience socioeconomic deprivation, community violence, maltreatment^7^ and lifestyle factors, it is conceivable that early environmental factors may alter the typical trajectory of brain maturation towards the end of childhood. Furthermore, we found that this accelerated brain age was most prominent when restricting the analysis to adolescents (ages 12–16) and, to a lesser extent, children (ages 6-11). These results highlight early- to mid- adolescence as a potential period during which brain aging appears most advanced in CD. Interestingly, this effect appears to be specific to those with a diagnosis of CD, as those with subthreshold CD failed to differ from controls. However, because the cohorts of youth with conduct problems without a formal CD diagnosis were younger (M = 11.7) than those in the main CD analysis (M = 13.4), this age difference may partly explain why the effect was not observed in this group. Nevertheless, these youth generally exhibit a less severe clinical profile, and deficits in cortical surface area and subcortical volume associated with CD did not generalize to this group in our previous study^16^.

### Callous-Unemotional Traits in CD Youth relate to Brain Age

Among youth with CD, it is estimated that up to 50% display high levels of callous-unemotional traits^3^. Hence, the DSM-5 included these traits (termed limited prosocial emotions) as a specifier for CD^1^. In the current study, we observed accelerated brain-PAD in youth with high callous-unemotional traits compared to TD youth, although they did not significantly differ from those with low CU traits. Prior work has demonstrated strong overlapping genetic influences between callous-unemotional traits and CD^38,39^ as well as more severe CD symptoms in youth with CD and high callous-unemotional traits (versus low callous-unemotional traits)^3,7^. Therefore, the stronger brain-PAD difference could reflect greater genetic loading. Yet, in youth with CD, brain-PAD did not correlated with severity of callous-unemotional traits and did not differ between CD subgroup with high and low callous-unemotional traits. One potential explanation is that callous-unemotional traits may interact with environmental factors in shaping maturational brain processes in youth with CD. While genetic factors explain most of the variance in the initial risk for early callous-unemotional traits, both genetic (distinct from initial risk) and non-shared environmental influences contribute significantly to the developmental course of callous-unemotional traits^40,41^. That said, CD youth with high and low callous-unemotional traits showed widely overlapping distributions of brain-PAD. This may partially be explained by null differences in cortical thickness between the two groups^16^, which appears to be more influential in the prediction of brain age, compared to other morphometric measures (see Supplementary Fig 3, see also^22^). Given the heterogeneous developmental trajectories of callous-unemotional traits^42,43^, the high/low cut-offs used in the current study may have obscured meaningful differences. Taken together, longitudinal studies tracking the co-development of CU traits and brain-PAD across multiple developmental stages could clarify the accelerated brain age observed in this study.

### Age-of-Onset of CD Youth relates to Brain Age

The DSM-5-TR (and earlier versions of the DSM) also accounts for the clinical heterogeneity of CD based on the age-of-onset of CD symptoms, i.e., whether they emerge before or after the age of 10^1^. This specifier was introduced following the seminal developmental taxonomic theory of antisocial behaviors^10^. In this theory, Moffitt^10^ posited two subgroups of individuals exhibiting antisocial behaviors, namely *life-course-persistent* and *adolescent-limited.* While the first subgroup was hypothesized to show a continuous and stable pattern of severe antisocial behaviors beginning before puberty (< 10 years old), the latter was hypothesized to show antisocial behaviors for a temporary period during adolescence. Despite showing superficially similar antisocial behaviors, Moffitt^44^ also argued that the underlying mechanism for engaging in antisocial behaviors may vary between the subgroups. Specifically, deficits in verbal and executive functions are believed to underlie early-onset antisocial behaviors, as demonstrated by poorer performance on neuropsychological tasks^10^. In contrast, the adolescent-onset subgroup is thought to engage in antisocial behaviors as an adaptive response to being trapped in a maturity gap between their biological and social age, a feature not found in the childhood-onset subgroup^44^. As such, we hypothesized that the childhood-onset CD subtype would exhibit decelerated brain age, reflecting its presumed neurodevelopmental origins, compared to the adolescent-onset CD subtype and typically developing youth. Instead, we found accelerated brain-PAD in youth with the adolescent-onset CD subtype compared to typically developing youth, although they did not significantly differ from those with childhood-onset CD. While no significant differences were observed between the two CD subgroups, the more pronounced difference in the adolescent-onset subgroup (relative to typically developing youth) is partially in line with Moffitt’s hypothesis. Indeed, she argued that youth following the adolescent-onset trajectory may engage in antisocial behaviors as an adaptive response to being caught in a maturity gap between their biological and social age, often mirroring the behavior of older peers.

Early adolescence is a critical developmental period marked by significant variability in pubertal growth^45^. Pubertal onset, rather than chronological age, may offer clearer insights into the different etiological pathways of CD. Indeed, the association between early pubertal timing and risk for disruptive behaviors^46^, and delinquency^47,48^ is well-documented. Furthermore, externalizing problems are among the strongest mental health correlates of the *puberty age gap:* the predicted pubertal age compared to chronological age^49^. Intriguingly, pubertal timing appears to moderate the heritability of CD. Specifically, the heritability of CD has been shown to be strongest (67%) in youth with average pubertal timing (around 12 years old), compared to only 8% for those with precocious puberty, suggesting a significant contribution of environmental influences on CD in the latter group^50^. These findings further support the idea that environmental factors -such as peer influences, parenting, and neighborhood factors -may ensnare those with early pubertal development into a persistent pattern of severe antisocial behaviors^48,51^. As environmental factors (e.g., abuse, maltreatment, exposure to violence, and neighborhood disadvantage) have also been linked to accelerated brain aging^36,37^, it could be argued that youth who engage in antisocial behaviors in adolescence may be particularly sensitive to socio-environmental influences during this critical developmental window, compared to those who start to engage in antisocial behaviors in childhood, who are presumably at greater genetic risk but do not show/experience such a maturity gap. While the current conceptualization defines CD subtypes based on chronological age (±10 years old), biological age (i.e., brain age, pubertal age) may offer a more sensitive approach for identifying inter-individual variability in the etiological pathways to CD.

### Strengths & Limitations

In this study, we report robust differences indicating accelerated brain age in youth with CD compared to typically developing peers. This effect was consistent across multiple international cohorts drawn from heterogeneous populations and across datasets with different acquisition protocols. Despite our rigorous analytic approach, several limitations must be acknowledged. First, the cross-sectional design prevented us from establishing whether accelerated brain age increases the risk for CD or, instead, the emergence of CD (and its related environmental features) accelerates maturational brain processes. Of note, despite a stronger effect found during adolescence, the developmental trajectory of brain-PAD among youth with CD remains largely unknown. As CD is characterized by different developmental pathways (e.g., childhood-onset, adolescent-onset), using prospective longitudinal designs may help clarify how brain-PAD maps onto those pathways over time (and based on our findings, whether those with a more advanced brain age experience the ‘maturity gap’ more keenly). Second, the BrainAGE model used in the current study showed somewhat variable performance across CD subtypes (Age-of-Onset, CU traits), suggesting that the findings should be interpreted with caution. It remains unclear whether these differences reflect true biological variation or are partly due to lower model performance and smaller sample sizes, as performance metrics are inherently linked to brain age (e.g., correlation between predicted and chronological age, mean absolute brain age difference). Third, the lack of consistency in assessment measures across cohorts prevented us from examining whether, and to what extent, accelerated brain age in youth with CD is associated with variations in pubertal development and/or socio-environmental risk factors (e.g., childhood adversity), and/or substance misuse, and/or clusters of CD symptoms. For instance, aggression and rule-breaking are known to be characterized by distinct developmental trajectories^52^ and neurobiological correlates^53^, leaving the possibility that these clusters of symptoms may be differentially associated with brain age. Because the brain-PAD difference between the CD and typically developing groups was most prominently found in adolescence, a period typically characterized by a sharp increase in rule-breaking behaviors (peak ≈15 years old)^52^, future studies should investigate the association between brain age and different CD symptom clusters. Fourth, while previous work has found that antisocial populations, including youth with CD, are characterized by greater variability in brain volume compared to controls^54^, our analyses demonstrated that a homoscedastic model provided a better fit to the data than a model with unequal residual variance (ΔBIC = 3.16). This suggested the residual variance was comparable between groups and that our main results were not driven by variance differences. Fifth, we acknowledge that our approach, based on traditional specifiers (e.g., differentiating between childhood- and adolescent-onset forms of CD, and high versus low CU traits), does not fully account for the substantial clinical and neurobiological heterogeneity observed in youth with CD, Future work -particularly using approaches such as normative modeling may allow for a finer-grained characterization of this heterogeneity, even after accounting for these so-called “*specifiers*”.

## Conclusion

This large collaborative effort, involving 14 international cohorts, identified a robust accelerated brain age of approximately five and a half months greater among youth with CD compared to typically developing youth. This effect appeared to be specific to early- to mid-adolescence and was stronger in those with high levels of callous-unemotional traits and those with adolescent-onset form of the disorder. Although these findings provide further support for conceptualizing CD as a neurodevelopmental disorder, prospective longitudinal studies are needed to better understand how this accelerated brain age unfolds over time and how it relates to variations in pubertal development and socio-environmental factors (e.g., childhood adversity and/or risky behaviors such as substance misuse).

## Methods

### Study samples

The current study pooled individual participant data from 14 (CD main analysis) and 10 (secondary analysis on elevated conduct problems) international cohorts within ENIGMA–ASB, comprising clinical, forensic, and community-based or population-based samples (for more details see^16^ and Supplementary Methods). Briefly, cohort eligibility criteria were a) a mean sample chronological age of 18 years or less, b) data available on sex, chronological age, and a diagnostic assessment of CD, and c) at least ten participants with CD or elevated conduct problems and 10 typically developing participants. More detailed information about inclusion criteria, subgrouping, and protocols can be found in Supplementary Methods and elsewhere^16^. Each contributing cohort had obtained ethical approval for their original study and for sharing de-identified data. This study was pre-registered (https://osf.io/u4d37) and the overall project received ethical approval from the University of Bath’s Psychology Research Ethics Committee (19-297/22-148). In the current study, a final sample of 1,119 youth with CD and 1,183 typically developing youth were included (see Supplementary Methods for more detailed information). Additionally, a total of 926 youth with elevated conduct problems and 922 typically developing youth^16^ were included in the secondary analysis testing the generalizability of the effect to subthreshold CD youth. This sample did not overlap with the CD sample from the main analysis. Demographic and clinical characteristics of the cohorts are presented in Supplementary Table 1.

### Image acquisition and pre-processing

Individual-level 3D T1-weighted volumetric brain magnetic resonance imaging data were pre-processed and quality controlled at the individual sites or project lead analysis sites (University of Birmingham and University of Bath) following standard ENIGMA imaging protocols^16^.

Neuroimaging data were pre-processed using FreeSurfer (version 5.3 or 6.0)^55^ and regions were parcellated based on the Desikan-Killiany and Aseg atlases.

We extracted global measures (i.e., total intracranial volume, average cortical thickness, and total surface area), as well as regional outcomes (i.e., cortical thickness and surface area for 68 cortical regions, and volume for 14 subcortical regions). The data were subsequently visually inspected, statistically evaluated for outliers (greater than 2.69 × SD), and pooled at the lead sites. For the current study, poorly segmented regions were excluded and imputed using the missForest R Package^56^, as used recently in the ENIGMA Consortium^57^. Participants with greater than 20% of missing neuroanatomical data were excluded [73 participants].

### Brain age prediction

The BrainAGE Global model of CentileBrain was used to predict brain age in the current mega-analysis (see https://centilebrain.org/)^58,59^. Twenty-one machine learning algorithms were initially trained on 2,105 typically developing children and adolescents (5-22 years old) from 5 cohorts and tested on two independent holdout datasets (n=4078, and n=594). The best performing algorithm using the Desikan–Killiany parcellation scheme was the Support Vector Regression with a Radial Basis Function ^58^. The model was then re-trained 35,683 healthy individuals (aged 5–90 years) and tested on an independent sample of 2102 healthy individuals, to examine the impact of harmonization strategies, chronological age range, and sample size on discovery and independent samples^59^. The best-performing model was retained, which included no cohort harmonization and two chronological age-bins (i.e., 5-40 and 40-90 years)^59^. In line with the BrainAGE Global framework, participant-level morphometric features were entered into the model separately for males and females. Brain-PAD was computed by subtracting chronological age from predicted neuroanatomical age. As in recent work ^18–20,22^, the model’s generalization performance was assessed through Pearson’s correlation and mean absolute error between predicted brain age and chronological age. A positive brain-PAD value indicates accelerated brain age compared to chronological age, while a negative value instead suggests a younger-appearing brain. Participants with brain-PAD beyond 1.5 times the interquartile range were considered outliers and excluded from further analyses [23 participants].

### Statistical Analyses

Given that previous work identified that the model estimating brain age without harmonization across cohorts best fit the data (BrainAGE Global^59^), we conducted linear mixed-effects models with random intercepts to estimate group differences in brain-PAD between children and adolescents with CD and typically developing, while accounting for the variability between cohorts. Chronological age and its quadratic term were included as covariates^60^ to mitigate the known bias whereby predicted age tends to be overestimated in younger populations^61^. BrainPAD of the *i-*th individual at *j-*th cohort was modeled as follows:

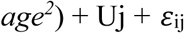

In the equation, intercept, diagnosis (CD), sex, chronological age, and chronological age^2^ effects were fixed. The terms U_j_ and ε_ij_ represent the random intercept attributed to the cohorts and the residual error, respectively. Cohen’s *d* was computed to enable comparisons with previous work that has examined brain age in other psychiatric disorders ^18–20,22^.

To assess the robustness of our findings across cohorts, we conducted jackknife resampling by systematically removing one cohort at a time and recalculating the beta estimate for each iteration. To account for possible overlap with healthy subjects from the ABCD cohort used in the BrainAGE Global model training, this additional analysis evaluated their impact on our overall findings. We subsequently tested for two- (diagnosis-by-chronological age; diagnosis-by-sex) and three-way (diagnosis-by-chronological age-by-sex) interactions. The main analysis was then repeated while covarying separately for comorbid attention-deficit/hyperactivity disorder, intelligence quotient, and medication status. Finally, we also explored whether brain-PAD was linearly associated with severity of conduct problems among youth with CD (see Supplementary Methods for more details).

Furthermore, to examine whether differences in brain-PAD between CD and typically developing participants would be moderated by developmental stages and/or sex, analyses were repeated by stratifying the samples with three chronological age bins approximating to the three stages of pubertal development^45^ (childhood [5-11 years old], adolescence [12-16 years old], late adolescence [17-21 years old]), and across sexes.

Subanalyses were conducted to identify sources of variation contributing to the brain-PAD difference. Specifically, we tested for potential differences between childhood-onset versus adolescent-onset CD subgroups (defined by symptom onset at age <10 years vs ≥10 years), and subgroups with low versus high callous-unemotional traits (defined by informant [self-reported or parent-reported], sex, and [for self-report] age-specific normative cutoffs on the Inventory of Callous-Unemotional traits (see Supplementary Methods for more details). We also tested whether callous-unemotional traits correlated with brain-PAD across the full sample, and specifically in youth with CD (see Supplementary Methods). Pairwise comparisons were adjusted for using the Benjamini-Hochberg false discovery rate method (*q*<0.05)^62^.

## Data Availability

The ENIGMA-Antisocial Behavior CD data used in the current study can be provided by the ENIGMA-Antisocial Behavior Working Group pending scientific review and a completed material transfer agreement. Request for the data should be submitted to: https://enigma.ini.usc.edu/ongoing/enigma-antisocial-behavior/.

## Code Availability

The analyses were conducted using BrainAGE Global model available online (https://centilebrain.org/).

## Acknowledgements

We thank all participants (and their families) for taking part in the studies that are included in this project. We also thank all researchers and support staff at the individual research sites for making this project possible by contributing their data. We gratefully acknowledge the late Prof. Shuqiao Yao for his outstanding contributions to conduct disorder research in China.

The ENIGMA consortium received funding from the National Institutes of Health (NIH) Consortium grant U54 EB020403, supported by a cross-NIH alliance that funds Big Data to Knowledge Centers of Excellence (BD2K).

Acknowledgments for the ENIGMA-Antisocial Behavior Working Group cohorts and contributors are as follows:

ABCD: Data used in the preparation of this article were obtained from the Adolescent Brain Cognitive Development^SM^ (ABCD) Study (https://abcdstudy.org), held in the NIMH Data Archive (NDA). This is a multisite, longitudinal study designed to recruit more than 10,000 children age 9-10 and follow them over 10 years into early adulthood. The ABCD Study® is supported by the National Institutes of Health and additional federal partners under award numbers U01DA041048, U01DA050989, U01DA051016, U01DA041022, U01DA051018, U01DA051037, U01DA050987, U01DA041174, U01DA041106, U01DA041117, U01DA041028, U01DA041134, U01DA050988, U01DA051039, U01DA041156, U01DA041025, U01DA041120, U01DA051038, U01DA041148, U01DA041093, U01DA041089,

U24DA041123, U24DA041147. A full list of supporters is available at https://abcdstudy.org/federal-partners.html. A listing of participating sites and a complete listing of the study investigators can be found at https://abcdstudy.org/consortium_members/. ABCD consortium investigators designed and implemented the study and/or provided data but did not necessarily participate in the analysis or writing of this report. This manuscript reflects the views of the authors and may not reflect the opinions or views of the NIH or ABCD consortium investigators. The ABCD data repository grows and changes over time. The ABCD data used in this report came from the <u>ABCD 3.0 data release</u>.

BESD was supported by a grant from the Netherlands Organization for Scientific Research–National Initiative Brain and Cognition (NWO-NIHC, project number 056-23-011), by a Leiden Institute Brain and Cognition starting grant to OFC, and an Örebro University grant to MA.

The Boys Town project was supported by Boys Town National Research Hospital and in part supported by the National Institute of Mental Health under award number K22-MH109558.

The Cambridge Female cohort was supported by the Wellcome Trust (grant numbers 069679 and 083140) and the Medical Research Council (MRC; grant code MC_US_A060_5PQ50).

The Cambridge Male cohort was funded by the Wellcome Trust (083140) and the MRC (grant code U.1055.02.001.00001.01).

CDKid was funded by a private donation; together with infrastructure support from the Mortimer D and Theresa Sackler Foundation, the MRC AIMS Network (G0400061/69344), an ongoing MRC-funded study of brain myelination in neurodevelopmental disorders (G0800298/87573), and the NIHR Biomedical Research Centre for Mental Health at King’s College London, Institute of Psychiatry and South London and Maudsley NHS Foundation Trust.

CSU-Yao was supported by grants from the National Nature Science Foundation of China (grant number 81471384).

CD-Zhou was supported by the National Natural Science Foundation of China (grant numbers 81571341, 62006038, and 82071543), Hunan Province Innovation Province Construction Project (2019SK2334), the Scientific Research Project of Sichuan Province Health Commission (20PJ213), the Natural Science Foundation of Hunan (2019JJ40424), Health Committee of Hunan (202103091470), and Clinical Medical Technology Innovation Guidance Project of Hunan (2020SK53415).

The FemNAT-CD consortium was funded by the European Commission under the 7th Framework Health Program (FP7/2007-2013, grant number 602407, coordinator CMF). This manuscript reflects only the authors’ views and the EU is not liable for any use that may be made of the information contained herein. AB was additionally supported by the Reiss Foundation Frankfurt am Main (grant numbers EER-2101-0002 and EER-2201–01).

Georgetown was supported in part by NIH/Eunice Kennedy Shriver National Institute of Child Health and Human Development (NICHD; to AAM, R03 HD 064906-01); The Georgetown-Howard Universities Center for Clinical and Translational Science (Ashley VanMeter NIH/NCATS 1KL2RR031974-01); Intellectual and Developmental Disabilities Research Center at Children’s National Medical Center (Ashley VanMeter, NIH/NICHD 2P30HD040677-11).

The IMAGEN consortium received support from the following sources: the EU-funded FP6 Integrated Project IMAGEN (Reinforcement-related behaviour in normal brain function and psychopathology; LSHM-CT-2007-037286), the Horizon 2020-funded ERC Advanced Grant STRATIFY (Brain network based stratification of reinforcement-related disorders; 695313), Horizon Europe environMENTAL (grant number 101057429), UK Research and Innovation (UKRI) Horizon Europe funding guarantee (10041392 and 10038599), Human Brain Project (HBP SGA 2, 785907, and HBP SGA 3, 945539), the Chinese Government via the Ministry of Science and Technology, The German Center for Mental Health (DZPG), the Bundesministerium für Bildung und Forschung (BMBF grants 01GS08152; 01EV0711; Forschungsnetz AERIAL 01EE1406A, 01EE1406B; Forschungsnetz IMAC-Mind 01GL1745B), the Deutsche Forschungsgemeinschaft (DFG grants SM 80/7-2, SFB 940, TRR 265, NE 1383/14-1), the Medical Research Foundation and MRC (grants MR/R00465X/1 and MR/S020306/1), the NIH-funded ENIGMA grants 5U54EB020403-05, 1R56AG058854-01, and U54 EB020403 as well as NIH R01DA049238, the National Institutes of Health Science Foundation Ireland (16/ERCD/3797), and the National Natural Science Foundation of China (grant 82150710554). Further support was provided by grants from: the French National Research Agency, ANR (ANR-12-SAMA-0004, AAPG2019 [GeBra]), the Eranet Neuron (AF12-NEUR0008–01 [WM2NA] and

ANR-18-NEUR00002–01 [ADORe]), the Fondation de France (00081242), the Fondation pour la Recherche Médicale (DPA20140629802), the Mission Interministérielle de Lutte-contre-les-Drogues-et-les-Conduites-Addictives (MILDECA), the Assistance-Publique-Hôpitaux-de-Paris and INSERM (interface grant), Paris Sud University IDEX 2012, the Fondation de l’Avenir (grant AP-RM-17-013), and the Fédération pour la Recherche sur le Cerveau, and the Ile-de-France Region (Action 16700103 -grant to QIM– VEAVE, n°23002745–23002747)

The KIND Lab Girls study was supported in part by a grant from the Hellman Fellows Program, a National Institute on Minority Health and Health Disparities sub-award (U54MD013368) from the UCR Center for Health Disparities Research, and a National Science Foundation CAREER award (NSF 2239067) awarded to KJM.

The MATRICS project has received funding from the EU’s Seventh Framework Programme for research, technological development, and demonstration under grant agreement number 603016. This manuscript reflects only the authors’ views and the EU is not liable for any use that may be made of the information contained herein.

The Aggressotype project has received funding from the EU’s Seventh Framework Programme for research, technological development, and demonstration under grant agreement number 602805, and from the Estonian Ministry of Education and Science project IUT20-40; Estonian Science Foundation grant 8622; European Regional Development Fund ERC Program TerVE (ELIKTU 3.2.10002.11-0002). This manuscript reflects only the authors’ views and the EU is not liable for any use that may be made of the information contained herein. BF gratefully acknowledges funding from the Netherlands Organization for Scientific Research (NWO) for the Growing up together in society (GUTS) project (grant number 024.005.011). NEH acknowledges additional support by the German Research Foundation (TRR379 projects B05 & Q01) and in parts by the Federal Ministry of Education and Research (Bundesministerium für Bildung und Forschung [BMBF]) and the Ministry of Science, Research and Arts of Baden-Württemberg within the German Center for Mental Health (DZPG, grants: DLR 01EE2504A, 01EE2504D).

MTwiNS was supported by grants from the National Institute of Mental Health (NIMH; no. R01-MH081813) and the Eunice Kennedy Shriver National Institute for Child Health and Human Development (NICHD; no. R01-HD066040) awarded to SAB, and by grants from the NIMH and the NICHD awarded to SAB and LWH (grant numbers UH3-MH114249 and R01-HD093334). Additional funding for data collection was provided by the Brain and Behavior Foundation (NARSADB Young Investigator Award to LWH), the Avielle Foundation (to LWH), and funds from the University of Michigan, as well as an NIH grant (S10OD026738) to support the MRI scanner. Any opinions, findings, conclusions, and recommendations expressed in this material are those of the authors and do not necessarily reflect the views of the NIH.

SAND was supported in part by the NIMH (R01-MH103761 to CSM and R01-MH121079 to LWH, CM, and CSM) and the NICHD (T32 HD007109-42), as well as an NIH grant (S10OD026738) to support the MRI scanner. SAND participants were recruited from the Future of Families and Child Wellbeing Study, which was supported by the NICHD under award numbers R01-HD036916, R01-HD039135, and R01-HD040421, as well as a consortium of private foundations. The content is solely the responsibility of the authors and does not necessarily represent the official views of the NIH.

The Southampton Family Study was funded by an Institute for Disorders of Impulse and Attention PhD studentship from the University of Southampton to KS and an Adventure in Research grant from the University of Southampton to GF.

UCL-T1/T2 was supported by a UK MRC grant (MR/K014080/1) and a UK Economic and Social Research Council grant (ES/N018850/1) to EV.

The Yale (Sukhodolsky) study was supported by grants from the NIMH (R01MH101514) and NICHD (R01HD083881) to DGS. KI is supported by the NIMH (K23MH128451), the fellowships on National Center for Advancing Translational Sciences grants (KL2 TR001862 and TL1 TR001864), and the Translational Developmental Neuroscience Training Program (T32 MH18268).

K23 was funded by the National Institute of Mental Health (NIMH; grant number K23-MH070036) and supported in part by grant number R01-MH080956 (both to MCS).

## Individual

AB was supported by the Federal Ministry of Research, Technology and Space (Bundesministerium für Forschung, Technologie und Raumfahrt; BMFTR) as part of the German Center for Child and Adolescent Health (DZKJ; code 01GL2405B).

CRKC is supported by the National Institutes of Health (Grant Nos. R21 MH139001; R01 MH129742).

HBW was supported by a Ruth L. Kirschstein National Research Service Award from the National Institute of Mental Health (F31 MH131373).

JRD was supported by a postdoctoral fellowship from the Canadian Institutes of Health Research (MFE-181885).

MS was supported by grant number ES/P000630/1 for the South West Doctoral Training Partnership awarded to the Universities of Bath, Bristol, Exeter, Plymouth, and West of England from the Economic and Social Research Council/UKRI. MS was also supported by UKRI under the UK government’s Horizon Europe / ERC Frontier Research Guarantee (BrainHealth, grant number EP/Y015037/1).

NJ was supported in part by NIH grant R01MH134004

YG was supported by the Newton International Fellowship funded by the Academy of Medical Sciences in the UK (grant agreement number NIF\R5\287).

NMR was supported by the Swiss National Science Foundation (105314_207624), the Hochschulmedizin Zurich (HMZ, STRESS), the University of Zurich Research Priority Program Adaptive Brain Circuits in Development and Learning (URPP AdaBD) and the Jacobs Foundation CRISP programme. GK is supported by a 2023 NARSAD Young Investigator Grant from the Brain & Behavior Research Foundation (30849).

MA was supported by The Netherlands Organization for Health Research and Development (ZonMw; research fellowship number 06360322210035), NWO (SSH Open Competition number 15810), Leiden University Fund (project Youth Mental Health Meets Big Data Analytics, grant number LUF23075-5-306), and Leiden University Fund (project grant number W213085-5).

SC was supported by the Biotechnology and Biological Sciences Research Council (BBSRC) and University of Birmingham funded Midlands Integrative Biosciences Training Partnership (MIBTP) (MIBTP2020: BB/T00746X/1)

DSP was supported by project ZIA-MH002781.

SADB is supported by an Economic and Social Research Council grant (ES/V003526/1).

## Author Information

### Authors and Affiliations

Accare Child Study Center, Groningen, The Netherlands

Andrea Dietrich, Pieter J. Hoekstra

ARA-BRAIN Institute II, Molecular Neuroscience and Neuroimaging, Forschungszentrum Jülich GmbH and RWTH Aachen, Juelich, Germany

Kerstin Konrad

Behavioural Science Institute, Radboud University, Nijmegen, The Netherlands

Maaike Oosterling

Birmingham Centre for Neurogenetics, School of Psychology, University of Birmingham, Birmingham, United Kingdom

Jack C. Rogers, Stephane A. De Brito

California Institute of Technology, Pasadena, CA, United States

Cindy C Hagan

Centre for Human Brain Health, School of Psychology, University of Birmingham, Birmingham, United Kingdom

Christopher D. Townsend, Nimrah Jabeen, Ruth Pauli, Sally C. Chester, Stephane A. De Brito, Yidian Gao

Centre for Developmental Science, School of Psychology, University of Birmingham, Birmingham, United Kingdom

Stephane A. De Brito

Centre for Population Neuroscience and Stratified Medicine (PONS), Department of Psychiatry and Neuroscience, Charité, University Medicine Berlin, Berlin, Germany and The Institute of Science and Technology for Brain-inspired Intelligence (ISTBI), Fudan University, Shanghai, China

Gunter Schumann

Child and Adolescent Mental Health Center, Copenhagen University Hospital and Mental Health Services CPH, Copenhagen, Denmark

Robert J.R. Blair

Child and Adolescent Psychiatry and Psychology Department, Hospital Clinic of Barcelona, Barcelona, Spain

Mireia Rosa-Justicia

Child Study Center, School of Medicine, Yale University, New Haven, CT, United States

Denis G. Sukhodolsky, Karim Ibrahim

Clinical Child Neuropsychology Section, Department of Child and Adolescent Psychiatry, Psychosomatics and Psychotherapy, University Hospital, RWTH Aachen, Aachen, Germany **Kerstin Konrad**

Cognitive Neuroscience, Freie Universität Berlin, Berlin, Germany

Leah E. Mycue

Department of Cancer Systems Imaging, Division of Diagnostic Imaging, The University of Texas MD Anderson Cancer Center, Houston, TX, United States

Sahil Bajaj

Department of Child and Adolescent Psychiatry and Psychology, 2021SGR01319, Hospital Clínic de Barcelona, IDIBAPS, CIBERSAM, Department of Medicine, Institute of Neuroscience, University of Barcelona, Barcelona, Spain

Josefina Castro-Fornieles

Department of Child and Adolescent Psychiatry and Psychotherapy, Central Institute of Mental Health, Medical Faculty Mannheim, Heidelberg University, Germany

Daniel Brandeis, Nathalie E. Holz, Pascal-M. Aggensteiner, Tobias Banaschewski

Department of Child and Adolescent Psychiatry and Psychotherapy, Psychiatric University Hospital, University of Zurich, Zurich, Switzerland

Daniel Brandeis

Department of Child and Adolescent Psychiatry, Faculty of Medicine, TU Dresden, Dresden, Germany

Anka Bernhard, Gregor Kohls

Department of Child and Adolescent Psychiatry, Institute of Psychiatry and Mental Health, Hospital General Universitario Gregorio Marañón, IiSGM, CIBERSAM, ISCIII, School of Medicine, Universidad Complutense, Madrid, Spain

Maria Jose Penzol Alonso

Department of Child and Adolescent Psychiatry, Psychosomatics and Psychotherapy, University Hospital Frankfurt am Main, Goethe University, Frankfurt am Main, Germany **Anne Martinelli, Christine M. Freitag**

Department of Child and Adolescent Psychiatry, University of Basel, Psychiatric University Hospital, Basel, Switzerland

Ana I. Cubillo, Christina Stadler

Department of Child and Adolescent Psychiatry and Psychology, Erasmus MC University Medical Center Rotterdam, Rotterdam, The Netherlands

Charlotte A.M. Cecil

Department of Child and Adolescent Psychiatry/Psychotherapy, University of Ulm, Ulm, Germany

Ulrike M.E. Schulze

Department of Clinical Medicine, Faculty of Health and Medical Sciences, University of Copenhagen, Denmark

Robert J.R. Blair

Department of Cognitive Neuroscience, Donders Institute for Brain, Cognition, and Behaviour, Radboud University Medical Center, Nijmegen, The Netherlands

Anouk H. Dykstra, Kim Lamers

Department of Education, Jianghan University, Wuhan, Hubei, China

Jibiao Zhang

Department of Epidemiology, Erasmus MC University Medical Center Rotterdam, Rotterdam, The Netherlands

Charlotte A.M. Cecil

Department of Forensic and Neurodevelopmental Sciences, The Institute of Psychiatry, Psychology and Neuroscience (IoPPN), King’s College London, London, United Kingdom **Arjun Sethi, Michael Craig**

Department of Human Genetics, Donders Institute for Brain, Cognition, and Behaviour, Radboud University Medical Center

Barbara Franke

Department of Medical Neuroscience, Donders Institute for Brain, Cognition, and Behaviour, Radboud University Medical Center, Nijmegen, The Netherlands

Barbara Franke, Jan K. Buitelaar

Department of Psychiatry, Amsterdam UMC, Location Vrije Universiteit Amsterdam, Amsterdam, The Netherlands

Moji Aghajani

Department of Psychiatry, Columbia University Irving Medical Center, New York, NY, United States

Dana E. Díaz

Department of Psychiatry, Leiden University Medical Center, Leiden, The Netherlands

Nic J.A. van der Wee

Department of Psychiatry, National Clinical Research Center for Mental Disorders, and National Center for Mental Disorders, The Second Xiangya Hospital of Central South University, Changsha, Hunan, China

Hui Chen, Jiansong Zhou, Xianliang Chen, Xiaoping Wang

Department of Psychiatry, The First Affiliated Hospital of Soochow University, Suzhou, Jiangsu, China

Qingsen Ming

Department of Psychiatry, Virginia Commonwealth University, Richmond, Virginia, United States

Robert J.R. Blair

Department of Psychology, Georgetown University, Washington, DC, United States

Abigail A. Marsh, Kathryn Berluti. Montana L. Ploe

Department of Psychology, Middlesex University London, London, United Kingdom

Karen Gonzalez-Madruga

Department of Psychology, School of Education Science, Hunan Normal University, Changsha, Hunan, China

Yali Jiang

Department of Psychology, The Catholic University of America, Washington, DC, United States

Elise M. Cardinale

Department of Psychology, University of Bath, Bath, United Kingdom

Alexandra Kypta-Vivanco, Graeme Fairchild, Harriet Cornwell, Marlene Staginnus, Rebecca L. Jackson

Department of Psychology, University of California, Riverside, Riverside, CA, United States

Kalina J. Michalska

Department of Psychology, University of Michigan, Ann Arbor, MI, United States

Christopher S. Monk, Heidi B. Westerman, Jules R. Dugré, Leah E. Mycue, Luke W. Hyde

Department of Psychology, Washington State University, Pullman, WA, United States

Montana L. Ploe

Department of Psychology, Yale University, New Haven, CT, United States

Arielle Baskin-Sommers, Karim Ibrahim

Department of Neurology, Max-Planck-Institute for Human Cognitive and Brain Sciences, Leipzig, Germany

Qiong Wu

Division of Psychology and Language Sciences, University College London, London, United Kingdom

Essi Viding, Harriet Phillips, Ruth Roberts

Faculty of Humanities, University of Utrecht, Utrecht, The Netherlands

Jilly Naaijen

German Center for Mental Health (DZPG), partner site Mannheim-Heidelberg-Ulm, Heidelberg, Germany

Pascal-M. Aggensteiner, Nathalie E. Holz

Hospital Universitario La Paz, IdiPAZ, School of Medicine, Universidad Autonóma de Madrid, CIBERSAM, Madrid, Spain

Celso Arango

Imaging Genetics Center, Mark and Mary Stevens Neuroimaging and Informatics Institute, Keck School of Medicine, University of Southern California, Marina del Rey, CA, United States

Christopher R.K. Ching, Melody J.Y. Kang, Neda Jahanshad, Paul M. Thompson, Sophia I. Thomopoulos

INSERM U1299 "Developmental Trajectories & Psychiatry", ENS Paris-Saclay, Mathematics center Borelli UMR 9010, University Paris-Saclay, Gif sur Yvette; & Research department, Barthelemy Hospital, Etampes; France

Jean-Luc Martinot

Institute for Mental Health, School of Psychology, University of Birmingham, Birmingham, United Kingdom

Jack C. Rogers, Sally C. Chester, Stephane A. De Brito

Institute for Social Research, University of Michigan, Ann Arbor, MI, United States

Colter Mitchell

Institute of Psychiatry, Psychology and Neurosciences, King’s College London, and the

Maudsley Hospital, London, United Kingdom

Paramala Santosh

Interdisciplinary Program in Neuroscience, Georgetown University, Washington, DC, United States

Abigail A. Marsh

Jacobs Center for Productive Youth Development at the University of Zurich, Department of Psychology, Zurich, Switzerland

Nora M. Raschle

Karakter Child and Adolescent University Center, Nijmegen, The Netherlands

Jan K. Buitelaar

Leiden Institute for Brain and Cognition, Leiden University Medical Center, Leiden, The Netherlands

Nic J.A. van der Wee

LUMC Curium, Child and Adolescent Psychiatry, Leiden University Medical Center, Leiden, The Netherlands

Charlotte P.S. Boateng, Robert Vermeiren

Medical Psychological Center, The Second Xiangya Hospital of Central South University, Changsha, Hunan, China

Daifeng Dong, Qingsen Ming, Ren Ma, Xiaoqiang Sun

Molecular Epidemiology, Department of Biomedical Data Sciences, Leiden University Medical Center, Leiden, The Netherlands

Charlotte A.M. Cecil

National Institute of Mental Health Intramural Research Program (NIMH-IRP), Bethesda, MD, United States

Daniel S. Pine

Neuroscience Center Zurich, University and ETH Zurich, Zurich, Switzerland

Nora M. Raschle

Olin Neuropsychiatry Research Center, Hartford, CT, United States

Michael C Stevens

Psychology School, Fresenius University of Applied Sciences, Frankfurt am Main, Germany

Anne Martinelli

Scaling, Research and Development, Child Development Institute, Toronto, ON, Canada

Areti Smaragdi

School of Academic Psychiatry, King’s College London, London, United Kingdom

Edmund J.S. Sonuga-Barke

School of Medicine, Conway Institute of Biomedical and Biomolecular Research, University College Dublin, Dublin, Ireland

Jeffrey C. Glennon

School of Psychology, The University of East Anglia, Norwich, United Kingdom

Yidian Gao

School of Psychology, University of Exeter, Exeter, United Kingdom

Ruth Pauli

School of Psychology, University of Southampton, Southampton, United Kingdom

Kate Sully

Section Forensic Family and Youth Care, Institute of Education and Child Studies, Leiden University, Leiden, The Netherlands

Moji Aghajani

Social, Genetic and Developmental Psychiatry Centre, Institute of Psychiatry, Psychology and Neuroscience (IoPPN), King’s College London, London, United Kingdom

Sylvane Desrivieres

Special Needs Education, Ghent University, Gent, Belgium

Olivier F. Colins

The Institute of Psychiatry, Psychology and Neuroscience (IoPPN), King’s College London, London, United Kingdom

Declan Murphy

The Institute of Psychiatry, Psychology and Neuroscience (IoPPN), King’s College London, London, United Kingdom

Ilyas Sagar-Ouriaghli

Trinity College Institute of Neuroscience and School of Medicine, Trinity College Dublin, Dublin, Ireland

Arun L.W. Bokde

Université Paris-Cité, Centre Borelli UMR 9010, Paris, France

Jean-Luc Martinot

University of Cambridge, Cambridge, UK

Luca Passamonti

University of Groningen, University Medical Center Groningen, Department of Child and Adolescent Psychiatry, Groningen, The Netherlands

Andrea Dietrich

University of Groningen, University Medical Center Groningen, Department of Psychiatry, Groningen, The Netherlands

Pieter J. Hoekstra

University of Rome “Tor Vergata”, Rome, Italy

Nicola Toschi, Silvia Minosse

Wu Tsai Institute, Yale University, New Haven CT, United States

Karim Ibrahim

ZfP Calw, Calw, Germany

Ulrike M.E. Schulze

## Contributions

**Conceptualisation:** AB-S, CAMC, DSP, GF, JRD, MS, MA, SADB, and YG.

**Analysis plan and methods:** AAM, AB-S, BF, CAMC, DSP, EV, GF, JKB, JRD, KJM, LWH, MS, MA, NJah, PMT, RJRB, SIT, SADB, and YG.

**Funding acquisition:** GF, JRD, MS, SADB, and YG (funding related to the individual cohorts is described in the Acknowledgments).

**Data acquisition, curation, and administration (individual cohorts, combined cohort, or both):** AAM, AIC, AD, AB, AB-S, AM, ASm, ASe, ALWB, BF, CA, CS, CMF, CSM, CCH, CM, DD, DED, DB, DM, DGS, EJSS-B, EDB, EMC, EV, GF, GK, GS, HCo, HP, HBW, HCh, IS-O, JC-F, JCR, JKB, J-LM, JCG, JianZ, JZha, JN, KJM, KG-M, KI, KS, KK, LP, LWH, MJPA, MS, MCS, MC, MR-J, MA, NEH, NJAvdW, NT, NMR, OFC, PJH, P-MA, QM, QW, RM, RJRB, RV, RP, RR, SAB, SB, SY, SADB, SD, TB, UMES, XC, XW, XS, YJ, and YG.

**Data analysis (individual cohorts):** AB, ASe, ALWB, CSM, CM, DED, DB, DGS, EMC, GS, HBW, JCR, JKB, JN, JC-F, KJM, KI, KB, LWH, MS, MCS, MR-J, MA, MLP, NJAvdW, NT, NMR, SB, SM, TB, XC, and YG.

**Quality control (individual cohorts):** AB, AK-V, AHD, ALWB, Ase, CPSB, CMF, CDT, DED, DGS, DM, EMC, EV, GS, HCo, HBW, JC-F, JCR, J-LM, JN, KJM, KI, KB, KL, LEM, MO, MS, MA, MLP, NJab, P-MA, RLJ, RP, SB, SCC, SM, SD, and YG.

**Statistical analysis (current study):** GF, JRD, MS, SADB, and YG.

**Interpretation of the findings:** AAM, AB, AB-S, BF, CAMC, CRKC, DM, DSP, DGS, EJSS-B, EV, GF, GK, JC-F, JKB, JRD, KJM, KI, LWH, MS, MJYK, MCS, MA, NJah, NMR, OFC, PMT, RJRB, SIT, SADB, SB, and YG.

**Writing of the original draft:** GF, JRD, MS, SADB, and YG.

**Review and editing:** AAM, AK-V, AIC, AD, AB, AB-S, AM, AHD, ASm, ASe, ALWB, BF, CA, CAMC, CPSB, CS, CMF, CDT, CRKC, CSM, CCH, CM, DD, DED, DB, DSP, DM, DGS, EJSS-B, EDB, EMC, EV, GF, GK, GS, HCo, HP, HBW, HCh, IS-O, JC-F, CR, JKB, J-LM, JCG, JianZ, JZha, JN, JRD, KJM, KG-M, KI, KS, KB, KK, KL, LEM, LP, LWH, MO, MJPA, MS, MJYK, MCS, MC, MR-J, MA, MLP, NEH, NJah, NJAvdW, NT, NJab, NMR, OFC, P-MA, PMT, PJH, QM, QW, RLJ, RM, RJRB, RV, RP, RR, SAB, SB, SCC, SM, SIT, SADB, SD, TB, UMES, XC, XW, XS, YJ, and YG.

*Due to data sharing and privacy regulations, not all authors had access to the full dataset but have granted full consent to the corresponding author to submit the current study for publication. GF, JRD, MS, SADB, and YG have accessed and verified the data. Authors are listed alphabetically by first name.

## Corresponding Authors

Correspondence to Jules R. Dugré & Stéphane A. De Brito

