## Supplementary Material for "Brain Age in Conduct Disorder: A Mega-Analysis of the ENIGMA Antisocial Behavior Working Group"

### Supplementary Methods

The inclusion criteria and (sub)grouping approach in the current study overlapped with that of a previous ENIGMA-ASB publication and hence, the below text overlaps with that presented in previous ENIGMA-ASB manuscripts.

#### 1. Grouping and subgrouping approach

While the specific inclusion/exclusion criteria differ by sample (please refer to [1]), the following exclusion criteria were applied for all analyses reported in this study: 1) a mean sample age > 18 years and individual participant age > 21 years, 2) IQ < 70 (where available) and 3) presence of neurological disorders, genetic syndromes, autism spectrum disorder as well as current diagnoses of schizophrenia or bipolar disorder (where the relevant information was available). We further excluded all participants without information on sex and age as these variables were included as covariates of no interest in the main analyses. Lastly, cohorts were only included if they comprised at least 10 participants with conduct disorder (CD) and 10 typically developing (TD) youth.

A note on autism spectrum disorder in the ABCD study (i.e., one of the included cohorts): A diagnosis of 'severe' autism spectrum disorder was an exclusion criterion within the ABCD study, but participants were retained in the ABCD study if parents/caregivers reported mild/moderate autism. Hence, it is possible that some participants with mild or moderate (parent-reported) autism spectrum disorder were retained in the ABCD subsample included in this study.

A note on the inclusion of participants over the age of 18 years: Although CD is most commonly diagnosed in childhood and adolescence, in the current study, we decided to include youths up to 21 years for the following reasons:

1. Consistency with prior ENIGMA research: Our study was designed to align with previous research conducted within the ENIGMA consortium, especially the work of Hoogman and colleagues [2,3] on attention deficit/hyperactivity disorder (ADHD), which adopted an age cut-off of 21 between adolescence and adulthood; as well as the work by Schmaal and colleagues [4] on major depressive disorder (MDD), which also set the age cut-off for adolescent versus adult analyses at ⩽21 years. This consistent/standardized approach was intended to ensure the comparability of results across disorders in youth, especially between CD and ADHD.

2. Continuity of brain development: Extensive longitudinal neuroimaging studies have demonstrated that brain maturation extends beyond adolescence, with significant structural changes observed until around age 21- 25 years. [5-7] This evidence supports the notion of emerging adulthood as a post-adolescent maturation stage, rather than a distinct phase after adolescence. Given this continuum of brain development and lack of biological justification to exclude individuals aged ≥ 18 from youth samples, we opted to include youths up to age 21 years.

3. Diagnostic criteria: According to the DSM-5-TR [8], an individual over 18 years old can still be diagnosed with CD if they do not meet the criteria for antisocial personality disorder. As there is no strong reason to exclude those aged above 18 from a psychiatric perspective, we opted to include them.

Based on the outlined reasons, we included youth up to age 21 years. However, to avoid shifting the mean age of the sample and maintain the focus on youth, we only included samples with a mean age of ≤ 18 years.

#### 2. Main Group Comparison (Conduct disorder versus Typically developing)

Participants allocated to the CD group had to have a current clinical or research diagnosis of CD. The former is provided in a healthcare setting by a medical professional, whereas the latter is primarily made for the purposes of a research study. For three of the 14 CD samples, it was not possible to fully verify the absence of these disorders in the control group as ADHD and/or ODD diagnoses were not provided. However, in each case the inclusion criteria indicated that controls were free of (current) psychiatric disorders and/or had no history of antisocial behaviours, indicating that these criteria were fulfilled by all samples.

#### 3. Subgroup comparisons

##### 3.1. Age-of-onset

In line with DSM-5-TR criteria [8], participants with CD who displayed CD symptoms before age ten were classified as having childhood-onset CD, whereas those with a later onset were classified as having adolescent-onset CD. Overall, information on age-of-onset was available for 722 CD participants (64.5% of the CD group) from seven out of 14 cohorts. Based on the described approach, 448 (62.0%) were allocated to the childhood-onset group and 274 (37.95%) were allocated to the adolescent-onset group. It should be noted that 57.14% of the childhood-onset participants were derived from the baseline release of the ABCD sample when participants were aged 9 or 10. TD youth from all cohorts were included in the control group.

##### 3.2. Low vs. high callous-unemotional (CU) traits

Data on Inventory of Callous-Unemotional Traits (ICU) was available in 9 out of 14 cohorts (Please refer to Table 1 in the Main Text). Youth with CD were divided into those high and low in CU traits based on the recently developed cut-offs by Kemp and colleagues [9] for the total scores of the ICU [10]. In contrast to the median split approach, which varies between studies, the use of normative cut-offs may facilitate replication in future analyses. In the current analyses, we applied the version-, gender-, and age-specific normative cut-offs, which have been shown to reduce the chance of false positives [9]. For the self-report version of the ICU, male youth with CD were allocated to the high-CU group if they scored ≥ 34 (up to age 14) or ≥ 37 (age 15 or older), while female youth with CD were allocated to the high-CU group if they scored ≥ 29 (up to age 14) or ≥ 32 (age 15 or older). For the parent-report version of the ICU, male youth with CD were allocated to the high-CU group if they scored ≥ 34 (any age), while female youth with CD were allocated to the high-CU group if they scored ≥ 30 (regardless of age). If participants had both self- and parent-report data available, they were included in the high-CU group if they were above the respective cut-off on either version. Overall, information on CU traits based on the ICU was available for 618 CD participants (55.23% of the whole CD group) from 9 out of 14 cohorts. More precisely, 263 (42.56%) were allocated to the low-CU group and 355 (57.44%) were allocated to the high-CU group.

#### 4. Severity of Conduct Disorder Symptoms

Information on CD symptoms was available for 800 CD participants (i.e., 506 with the CD-oriented subscale of the Child Behavior Checklist [11], and 326 with the CP subscale of the Strengths and Difficulties Questionnaire [12]). To harmonize scores between the two measures, the percentage of maximum possible score was computed for each participant. If a participant was assessed using both measures, DSM-Oriented CBCL score was chosen given its correspondence to DSM-5 criteria.

#### 5. Outliers Detection & Removal

Across the 14 cohorts on CD, seventy-three participants (73) were excluded from the CentileBrain BrainAGE2 prediction model due to having >20% missing neuromorphometric data. In addition, twenty-three individuals (23) showed greater brain-PAD beyond 1.5 times the IQR (14 CD, 8 TD) and were therefore excluded from further analyses.

Across the 10 cohorts contributing youth with elevated CP, twenty-five participants (25) were discarded from the CentileBrain BrainAGE2 prediction model due to having >20% missing neuromorphometric data. In addition, one hundred and four individuals (104) were considered outliers (62 CP, 42 TD), and were therefore excluded from further analyses.

**Supplementary Table 1.** Demographic and clinical characteristics of the included cohorts in the Replication Analysis

| Cohorts | Total N | Typically developing youth | | | | Elevated Conduct Problems | | | |
|---|---|---|---|---|---|---|---|---|---|
| | | n | F:M | Age, yrs | IQ | n | F:M | Age, yrs | IQ |
| ABCD (3.0, baseline) [a,b] | 810 | 407 | 192:215 | 9.5 (0.50) | 96.0 (15.38) | 403 | 184:219 | 9.5 (0.50) | 95.5 (15.51) |
| FemNAT-CD [a,b] | 23 | 13 | 6:7 | 13.5 (2.73) | 108.3 (13.67) | 10 | 6::4 | 13.4 (2.91) | 100.8 (13.40) |
| Georgetown | 68 | 37 | 20:17 | 13.1 (2.34) | 111.7 (13.9) | 31 | 13:18 | 14.0 (2.37) | 98.3 (10.19) |
| IMAGEN (baseline) [a,b] | 669 | 334 | 163:171 | 13.9 (0.48) | - | 335 | 167:168 | 14.0 (0.47) | - |
| KIND Lab girls study | 39 | 28 | 28:0 | 9.54 (1.23) | 96.3 (9.94) | 11 | 11:0 | 10.2 (1.17) | 95.9 (8.24) |
| MATRICS/Aggressotype [a,b] | 57 | 31 | 2:29 | 12.5 (2.47) | 102.7 (11.51) | 26 | 2:24 | 12.6 (2.79) | 100.1 (11.23) |
| MTwiNS [a,b] | 10 | 8 | 7:1 | 13.4 (2.39) | 108.0 (0.00) | 2 | 0:2 | 13.5 (2.12) | 95.0 (11.31) |
| SAND [b] | 17 | 8 | 2:6 | 15.3 (0.46) | - | 9 | 1:8 | 15.3 (0.50) | - |
| UCL-T1/T2 | 55 | 14 | 0:14 | 13.65 (1.65) | 98.8 (8.98) | 41 | 0:41 | 14.0 (1.40) | 98.5 (12.68) |
| Yale [b] | 100 | 42 | 15:27 | 11.6 (1.86) | 112.1 (13.56) | 58 | 21:37 | 10.7 (1.84) | 107.3 (14.66) |
| Total (11 samples) | 1848 | 922 | 435:487 | 11.6 (2.34) | 98.9 (15.63) | 926 | 405:521 | 11.7 (2.38) | 97.3 (15.07) |

The reported values reflect n or mean (+ standard deviation). Information on sex and age were available for all participants, whereas IQ was not available for all samples or all participants within a sample.

[a] Multi-site/-scanner sample.

[b] Control group matched on age and sex (and IQ, if available) using propensity score matching.

F:M=female:male participant sex ratio. IQ=intelligence quotient.

**Supplementary Table 2.** Chronological Age, Brain Age, and Brain Predicted Age Differences per cohorts from the **Conduct Problems Analyses**

| Cohorts | Controls | | | | Youth with Conduct Problems | | | |
|---|---|---|---|---|---|---|---|---|
| | **n** | **Chronological age** | **Brain Age** | **Brain-PAD** | **n** | **Chronological age** | **Brain Age** | **Brain-PAD** |
| ABCD (3.0, baseline) [a,b] | 407 | 9.5 (0.50) | 8.3 (1.52) | -1.18 (1.44) | 403 | 9.5 (0.50) | 8.64 (1.75) | -0.84 (1.70) |
| FemNAT-CD [a,b] | 13 | 13.5 (2.73) | 14.0 (4.04) | 0.46 (1.90) | 10 | 13.4 (2.91) | 12.6 (3.25) | -0.84 (1.03) |
| Georgetown | 37 | 13.1 (2.34) | 13.0 (3.42) | -0.17 (2.13) | 31 | 14.0 (2.37) | 13.4 (3.64) | -0.63 (2.43) |
| IMAGEN (baseline) [a,b] | 334 | 13.9 (0.48) | 13.9 (1.13) | -0.07 (1.12) | 335 | 14.0 (0.47) | 13.9 (1.19) | -0.09 (1.16) |
| KIND Lab girls study | 28 | 9.54 (1.23) | 8.82 (1.91) | -0.71 (1.81) | 11 | 10.2 (1.17) | 10.0 (2.39) | -0.18 (1.97) |
| MATRICS/Aggressotype [a,b] | 31 | 12.5 (2.47) | 11.6 (3.83) | -0.85 (2.51) | 26 | 12.6 (2.79) | 12.5 (2.69) | -0.08 (2.47) |
| MTwiNS [a,b] | 8 | 13.4 (2.39) | 13.9 (3.44) | 0.52 (1.57) | 2 | 13.5 (2.12) | 14.4 (0.58) | 0.86 (1.54) |
| SAND [b] | 8 | 15.3 (0.46) | 15.8 (2.46) | 0.53 (2.27) | 9 | 15.3 (0.50) | 14.81 (2.02) | -0.52 (2.22) |
| UCL-T1/T2 | 14 | 13.65 (1.65) | 14.0 (3.14) | 0.53 (2.06) | 41 | 14.0 (1.40) | 13.8 (2.89) | -0.24 (2.21) |
| Yale [b] | 42 | 11.6 (1.86) | 11.5 (2.55) | -0.13 (1.82) | 58 | 10.7 (1.84) | 9.89 (2.60) | -0.80 (1.98) |
| Total (11 samples) | 922 | 11.6 (2.34) | 11.1 (3.22) | -0.59 (1.58) | 926 | 11.7 (2.38) | 11.2 (3.11) | -0.50 (1.67) |

*Note.* Brain Age and Brain-PAD were computed using BrainAGE2 model of CentileBrain. Brain Age refers to Neuroimaging-Predicted Age. Brain-PAD = Brain-Predicted Age Difference (Δ brain age - chronological age)

[a] Multi-site/-scanner sample.

[b] Control group matched on age and sex (and IQ, if available) using propensity score matching.

F:M=female:male participant sex ratio. IQ=intelligence quotient.

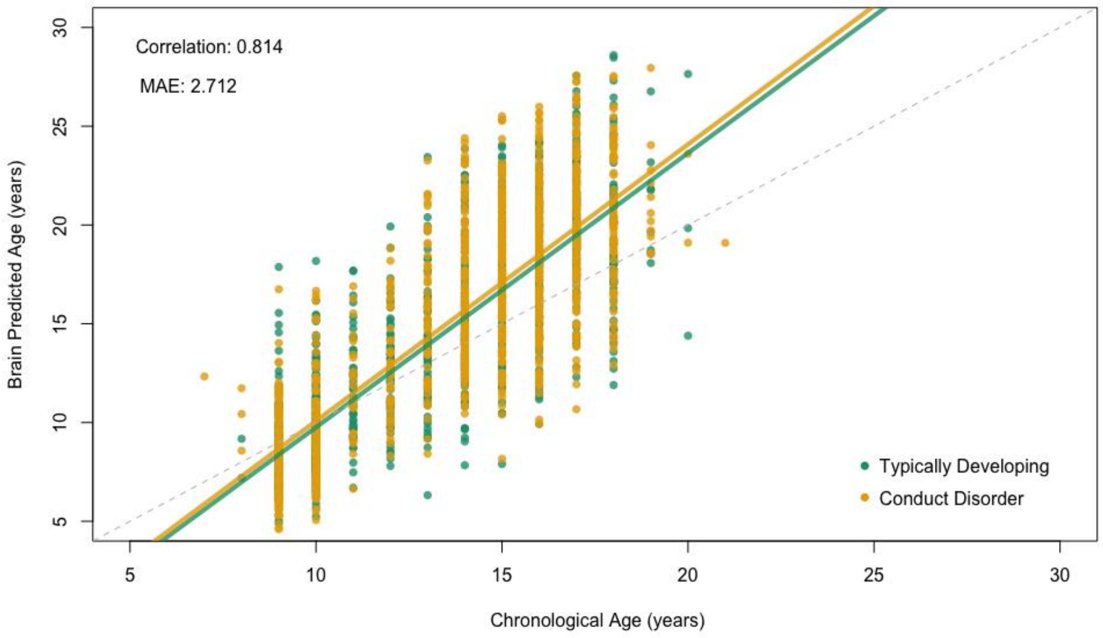

**Supplementary Fig. 1.** Centile Brain's BrainAGE2 model generalization across youth with CD and TD from 14 international cohorts (r = 0.81, MAE = 2.71 years). The model generalized similarly well across CD ($r$ = 0.81, MAE = 2.88) and TD ($r$ = 0.82, MAE = 2.56) and across $CD_{females}$ (r = 0.80, MAE = 2.82), $CD_{males}$ (r = 0.81, MAE = 2.90), $TD_{females}$ (r = 0.80, MAE = 2.56), and $TD_{males}$ (r = 0.83, MAE = 2.56).

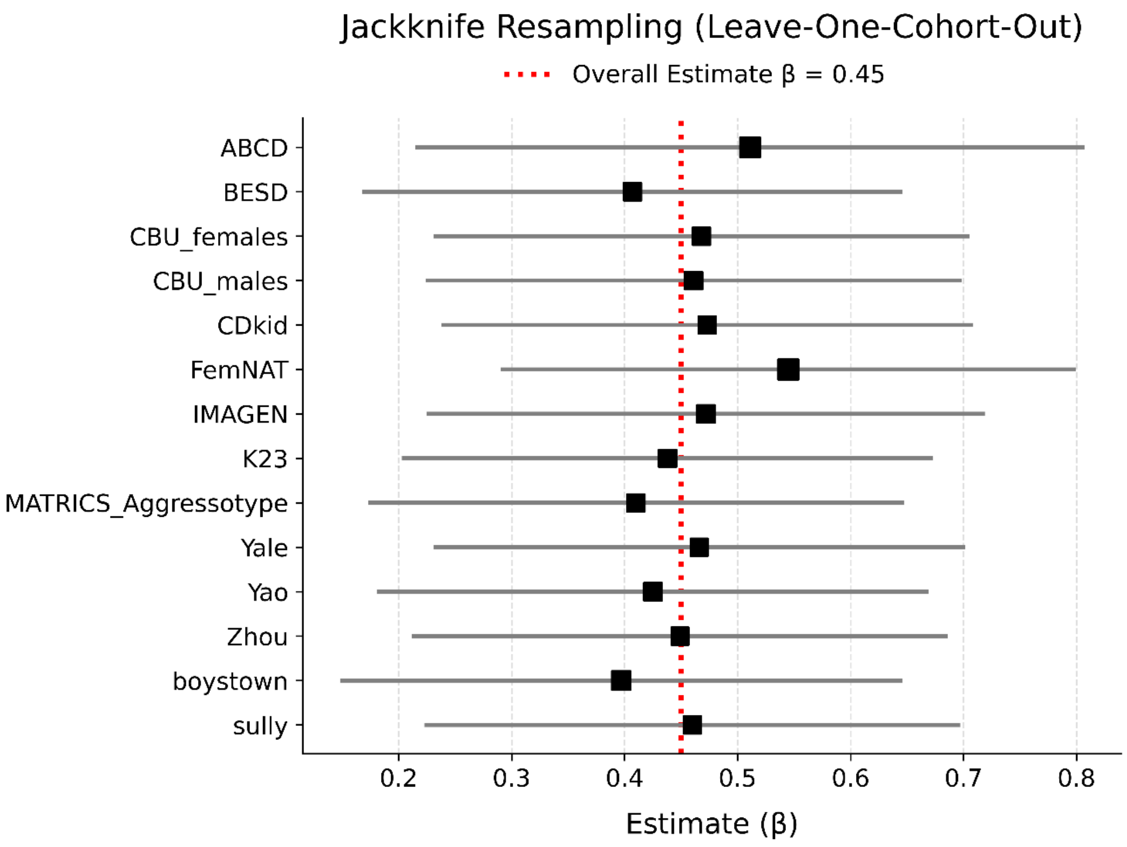

**Supplementary Fig. 2.** Results of the Jackknife resampling analysis. Forest plot shows BrainAGE estimates after removing each cohort sequentially. Each square is proportional to the cohort size.

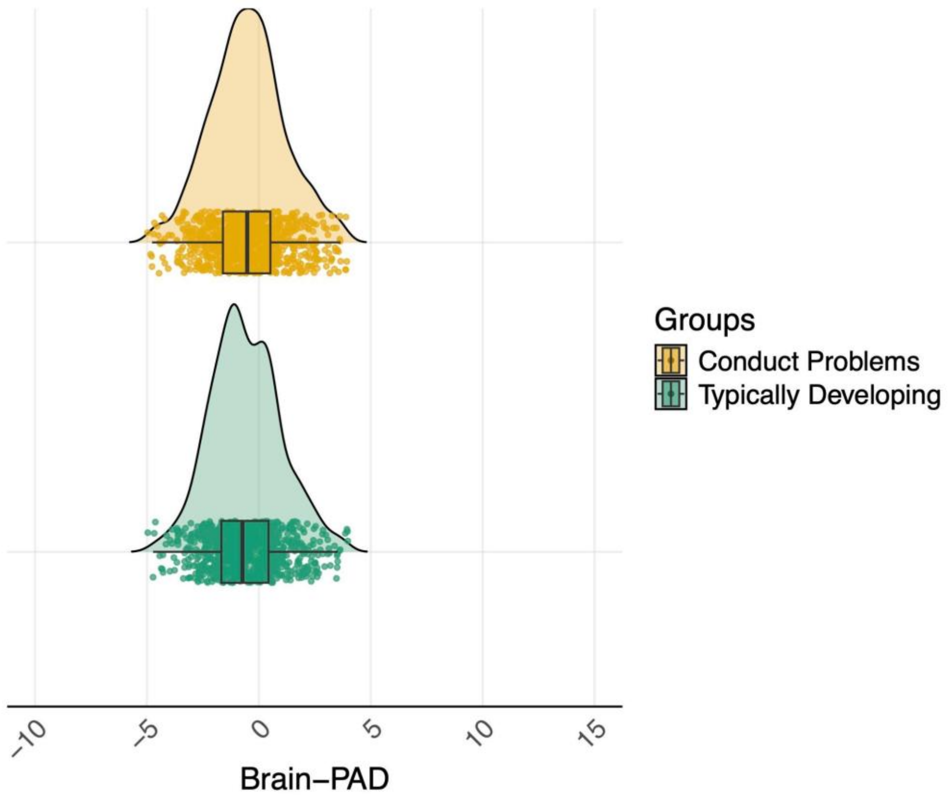

**Supplementary Fig. 3.** Absence of significant difference in brain aging between youth with elevated conduct problems (n=926) and typically developing youth (n=922).

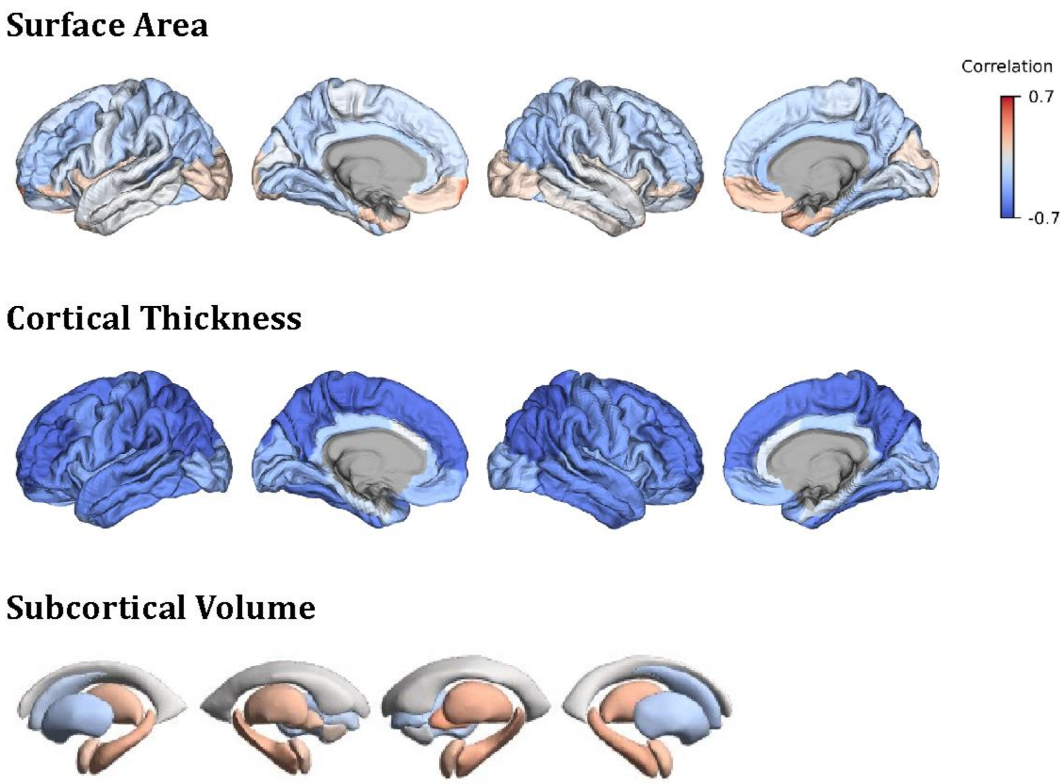

**Supplementary Fig. 4.** Correlation Coefficients of brain age and FreeSurfer's morphometric measures across in youth with CD and TD.
